# Cryo-EM reveals the structure and dynamics of a 723-residue malate synthase G

**DOI:** 10.1101/2022.08.19.504428

**Authors:** Meng-Ru Ho, Yi-Ming Wu, Yen-Chen Lu, Tzu-Ping Ko, Kuen-Phon Wu

**Affiliations:** Institute of Biological Chemistry, Academia Sinica, Taipei, Taiwan, 115; Institute of Biochemical Science, National Taiwan University, Taipei, Taiwan, 106

## Abstract

Determination of sub-100 kilodalton (kDa) structures by cryo-electron microscopy (EM) is a longstanding but not straightforward goal. Here, we present a 2.9-Å cryo-EM structure of a 723-amino acid apo-form malate synthase G (MSG) from *Escherichia coli*. The cryo-EM structure of the 82-kDa MSG exhibits the same global folding as structures resolved by crystallography and nuclear magnetic resonance (NMR) spectroscopy, and the crystal and cryo-EM structures are indistinguishable. Analyses of MSG dynamics reveal consistent conformational flexibilities among the three structural approaches, most notably that the α/β domain exhibits heterogeneity. We observed that sidechains of F453, L454, M629, and E630 residues involved in hosting the cofactor acetyl-CoA and substrate rotate differently between the cryo-EM apo-form and complex crystal structures. Our work demonstrates that the cryo-EM technique can be used to determine structures and conformational heterogeneity of sub-100 kDa biomolecules to a quality as high as that obtained from crystallography and NMR spectroscopy.

## Introduction

Recent advancements in cryo-EM techniques have established that this methodology, alongside X-ray crystallography and NMR, is a practical experimental tool for determining three-dimensional biomolecular structures ^1–4^. However, cryo-EM and NMR represent techniques at the two ends of a spectrum in terms of target molecular sizes. For example, as of August 2022, < 50 NMR-derived protein structures of molecular weight (MW) >50 kDa had been deposited in the Protein Data Bank (PDB). Since the often crowded and overlapping spectral cross-peaks of large MW proteins impede atomic spin assignment, spectral and structural determination of large proteins by NMR requires selectively labeled samples and multiple experiments, resulting in a time-consuming analysis ^5–7^. In contrast, cryo-EM involves analyzing particles imaged by direct electron detectors, requiring distinguishable and sufficient contrast of objects in a noisy background. Therefore, cryo-EM tends to be deployed for 200-500 kDa proteins or complexes, and rarely for those of sub-100 kDa.

Nevertheless, pioneering studies have demonstrated the feasibility of applying cryo-EM to determine sub-100 kDa protein structures, including 3.2 Å hemoglobin (64 kDa) ^8^, 2.7 Å horse live alcohol dehydrogenase (82 kDa, C2 dimer) ^9^, 2.8 Å human methemoglobin (64 kDa, C4 tetramer) ^9^, 3.2 Å streptavidin (52 kDa, D2 tetramer) ^10^, and 2.8 Å human CDK-activating kinase (CAK) complexed with cyclin H (84 kDa, heterodimer) ^11^. These proteins all possess small particle sizes and, apart from CAK-cyclin H, they are symmetric assemblies. Sample preparation, data collection, processing, and single particle analysis for cryo-EM of 52-100 kDa proteins necessitate strategies distinct from those of mega-Dalton ribosomes ^12^ and proteasomes ^13^. However, further refinements of procedures for sub-100 kDa cryo-EM are needed to enhance its applicability.

Here, we present a cryo-EM structural analysis of a 723-amino acid (aa) *E. coli* malate synthase G (ecMSG). Bacteria possess two isoforms of malate synthase (A or G) in the glyoxylate cycle. MSG uniquely bypasses a few steps of the citric acid cycle ^14,15^ to produce malate and coenzyme A (CoA) by consuming glyoxylate and acetyl-CoA. MSG has been proposed as a drug target for the virulence of opportunistic pathogens such as mycobacteria ^16,17^ and *Pseudomonas aeruginosa* ^18^. Moreover, structures of 82-kDa MSG have previously been determined by X-ray crystallography (1D8C ^19^, 1P7T ^20^, both representing ligand-bound structures) and solution NMR spectroscopy ^21,22^. The cryo-EM structures presented herein enable structural comparison of a single protein using three primary tools of structural biology.

## Results and Discussion

### The cryo-EM and crystal structures of apo-form ecMSG

We amplified the *ecMSG* gene directly from the DH5α *E. coli* strain, before conducting affinity and size-exclusion purification steps. To investigate the status of the molecular assembly, purified ecMSG was subjected to size-exclusion chromatography combined with multi-angle light scattering (SEC-MALS) (Figure 1A), which indicated an average MW of 82.6 kDa (theoretical MW = 82.9 kDa with a His-tag) and a hydrodynamic radius of 32.1 Å, confirming its monomeric state in solution. Then we prepared an apo-form ecMSG (substrate- and Mg^2+^-free) for cryo-EM data collection and structural determination. The grids containing 0.1-0.5 mg/ml ecMSG revealed that sufficient amounts of apo-form particles had been acquired by our K2 direct detector on a 300 kV Titan Krios microscope (Figure 1B). 2D classifications indicated particle sizes of ~70 Å, with a shape and structural features typical of ecMSG (Figure 1C). We selected 211,582 particles from 700 micrographs for 3D reconstruction using a non-uniform refinement approach in cryoSPARC 3.2 ^23^, resulting in a 2.9 Å map. This map vividly displays the secondary structural regions and residue sidechains of ecMSG (Figure 1D-E). Particularly well characterized are the most extended α-helix (comprising residues A32-S70), anti-parallel β-sheets, aromatic residues F35 and W67, and long sidechain residues L46, R57, K232, and I242 (Figure 1F). The local resolution estimate of this 2.9-Å map illustrates that most residues are in the range of 2.4-2.8 Å, except for residues E151-V155, Q299-K312, and the N/C-termini (Figure 1G). Notably, residues Q299-K312 in previous crystal structures (1D8C, 1P7T) are not well resolved, implying a highly dynamic β-hairpin region. Our cryo-EM data indicate that the 82-kDa ecMSG 3D structure has been successfully determined.

**Figure 1.**
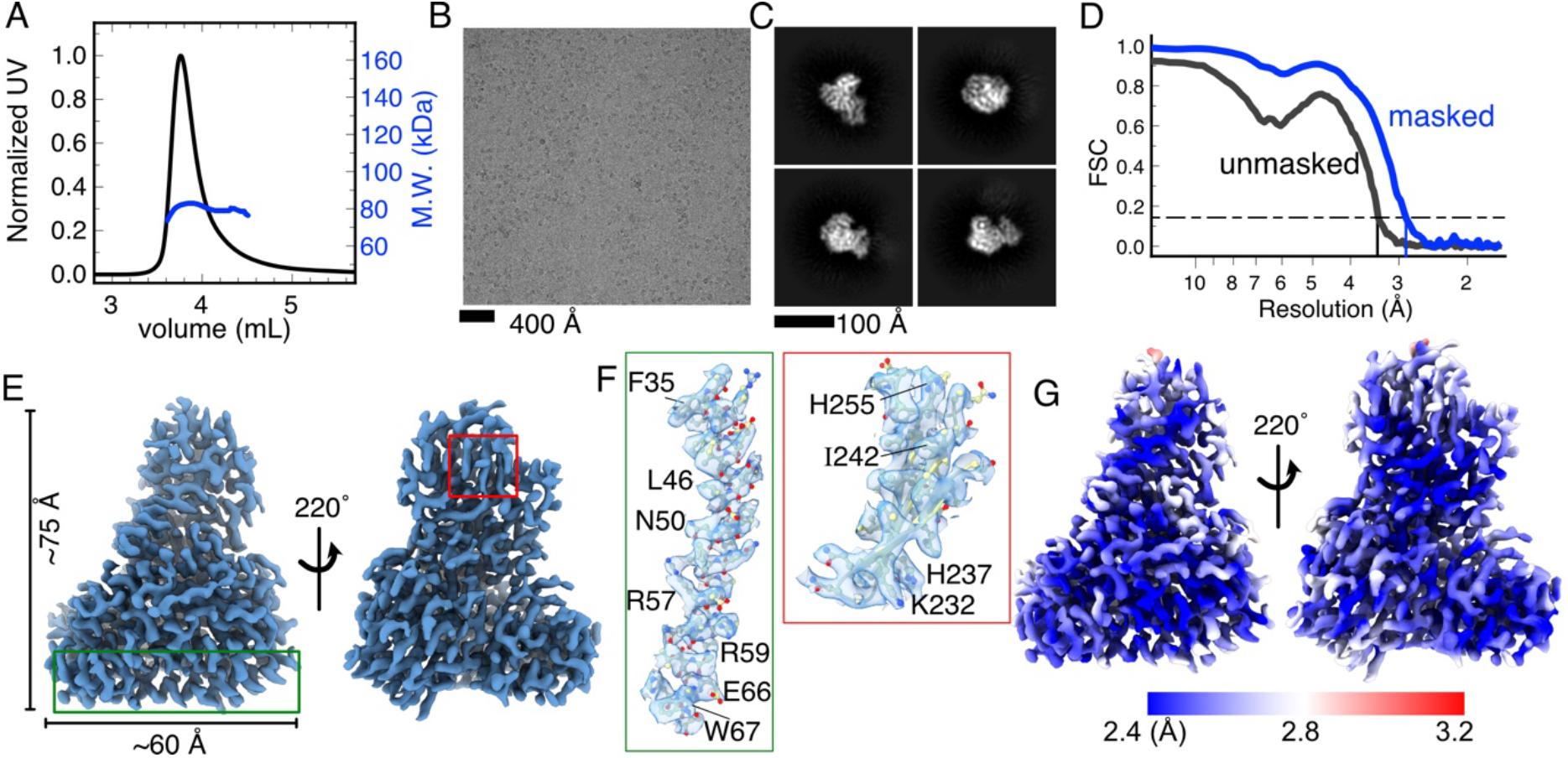
The cryo-EM data of ecMSG. (A) A SEC-MALS experiment reveals that ecMSG is typically monomeric, with an average mass of 82.61 kDa. (B) Representative micrograph of ecMSG, revealing good contrast and density of MSG particles. (C) Representative 2D classifications display different orientations, but consistent sizes of ecMSG. (D) The reconstructed 2.9-Å map of ecMSG. (E) The 3D map of ecMSG generated using ChimeraX 1.4 ^41^ (contour level: 0.35) reveals the characteristics of many helices and strands. (F) Boxed areas in (E) shown in map reveal clear structural features and sidechains of selected antiparallel β-sheets and the longest α-helix (residues 33-70). Long, aromatic or charged sidechains have been labeled to reveal the clear cryo-EM map densities of the sidechains. (G) The estimated local resolution of the ecMSG cryo-EM map covers a resolution range of 2.4-3.2 Å.

We built the ecMSG structure based on this 2.9-Å map, revealing a virtually identical 3D structure to substrate-bound MSG crystal structures (PDB ID: 1P7T, 1D8C). We wanted to directly compare the structures of ecMSG determined by crystallography ^19,20^, solution NMR spectroscopy ^21,22^, and cryo-EM (this work). However, the previously resolved ecMSG crystal structures were glyoxylate- or pyruvate:acetyl-CoA-bound (1D8C and 1P7T, respectively). In contrast, the cryo-EM and NMR ecMSG structures are cofactor- and substrate-free. Moreover, we identified 26 and 14 incomplete ecMSG sidechains in the 1P7T and 1D8C complex structures, respectively. Therefore, we determined a new crystal structure of apo-form ecMSG at 1.6 Å to enable a comparative analysis of ecMSG structures under consistent conditions.

### Structural comparisons of apo-form ecMSG

The apo-ecMSG structures determined by cryo-EM (2.9 Å), crystallography (1.6 Å, two versions), and an NMR ensemble (10 conformers) ^22^ are shown in Figure 2A, in which only the first NMR conformer is used. The apo-ecMSG crystal structure displays a dimer in one asymmetric unit presenting clear and preserved sidechains, except for unresolved residues E303-R307 in a β-hairpin. The root mean square deviations (RMSDs) of the globally aligned apo-ecMSG structures are < 0.7 Å, implying that the cryo-EM and crystal structures are remarkably indistinguishable. Our detailed analysis of secondary structures uncovered that the cryo-EM structure is composed of 24 α-helices, 8 3_10_-helices, and 26 β-sheets, whereas the 1.6-Å crystal structure (using chain A) has 24 α-helices, 11 3_10_-helices, and 26 β-sheets (Figure 3). The NMR-derived ecMSG structure exhibits the same global fold ^22^ as both the cyro-EM and crystal structures, but 9 β-sheets in the α/β domain were not well resolved in the NMR ensemble (Figure 3), resulting in a large RMSD of 4 Å between the NMR and cryo-EM ecMSG structures (Figure 2B). Moreover, helices encompassing residues 589-613 and 653-671 of the ecMSG NMR structure are shifted toward the α/β domain by 12 Å relative to both the cryo-EM and crystal structures (Figure 2C). Since the NMR data were acquired at 37 °C, heat could be a factor contributing to a more dynamic conformation compared to the frozen conditions under which the crystal and cryo-EM structures were acquired.

**Figure 2.**
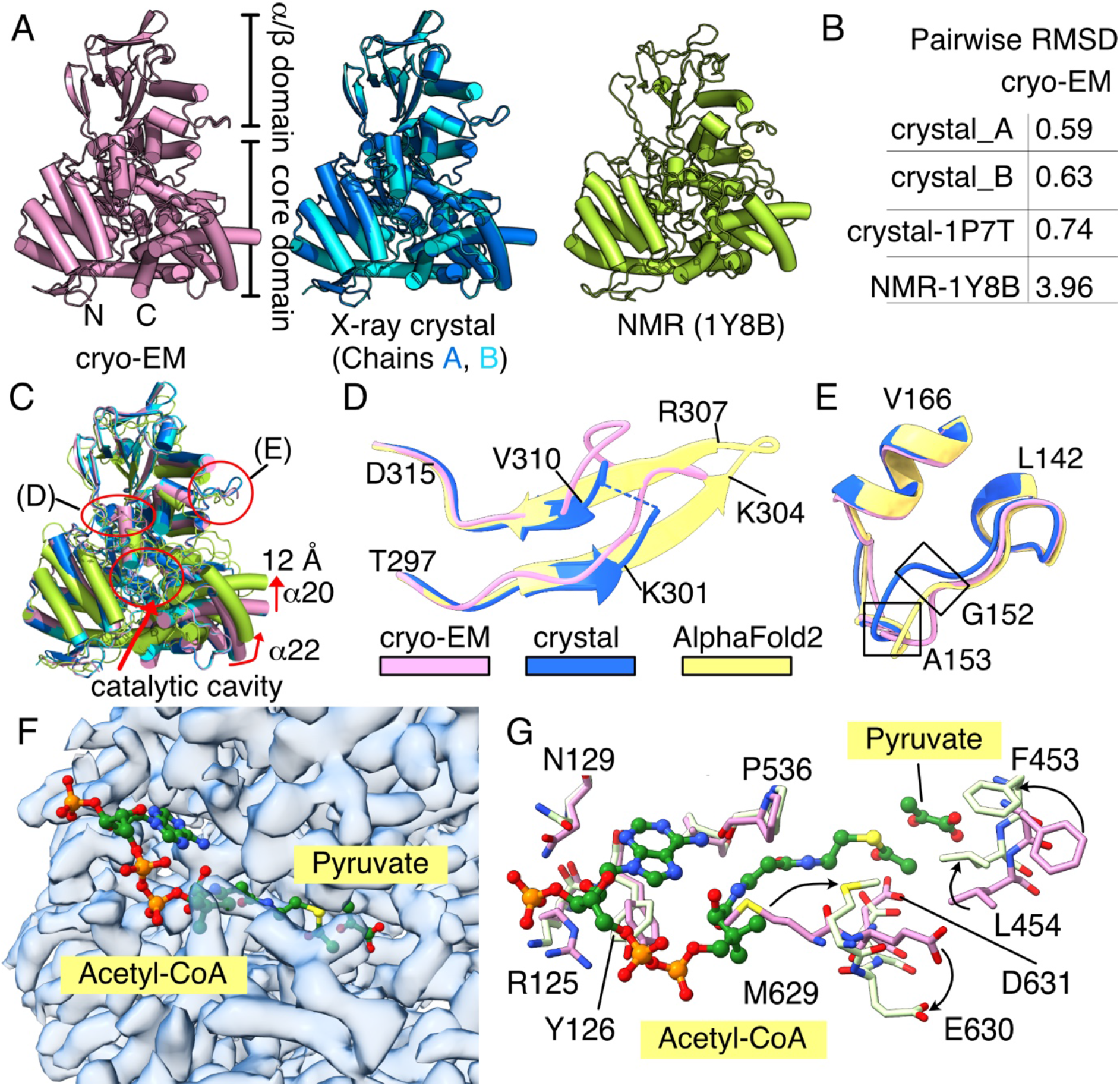
Comparison of apo-form ecMSG structures. (A) Alignments and superimpositions of apo-form ecMSG structures determined by cryo-EM (pink), X-ray crystallography (cyan and blue), and solution NMR spectroscopy (green). Globally, protein folding is similar amongst methodologies, but the NMR structure ensemble is more dynamic. (B) Calculated RMSDs (Å) of global structural alignments paired to the cryo-EM ecMSG structure. (C) The aligned cryo-EM, crystal, and NMR structures of apo-ecMSG illustrate that the α20 and α22 helices of the NMR structure are shifted ~12 Å toward the α/β-domain, whereas the cryo-EM and crystal structures are identical. Circles indicate variations analyzed in panels D, E and F. (D) Localized structure of ecMSG residues 297-315 are distinctive among the three different methodologies, including Alphafold2 (yellow) ^26^, suggesting a dynamic feature. (E) A loop (residues 142-160) between helices α6 and α7 is also dynamic. Residues G152 and A153 are labeled to highlight divergence among the three structures. (F) Aligned acetyl-CoA and pyruvate from the crystal structure (1P7T) reveals an unoccupied space in the catalytic pocket of the apo-form cryo-EM map. (G) Sidechain comparisons of the apo- (cryo-EM, pink) and complex-form ecMSG (1P7T, light green) structures illustrate that residues F453 and L454 flip and stably clamp the pyruvate (arrows). Residues M629 and E630 are oriented outward to provide space for hosting the carbon chain of acetyl-CoA (arrow).

**Figure 3.**
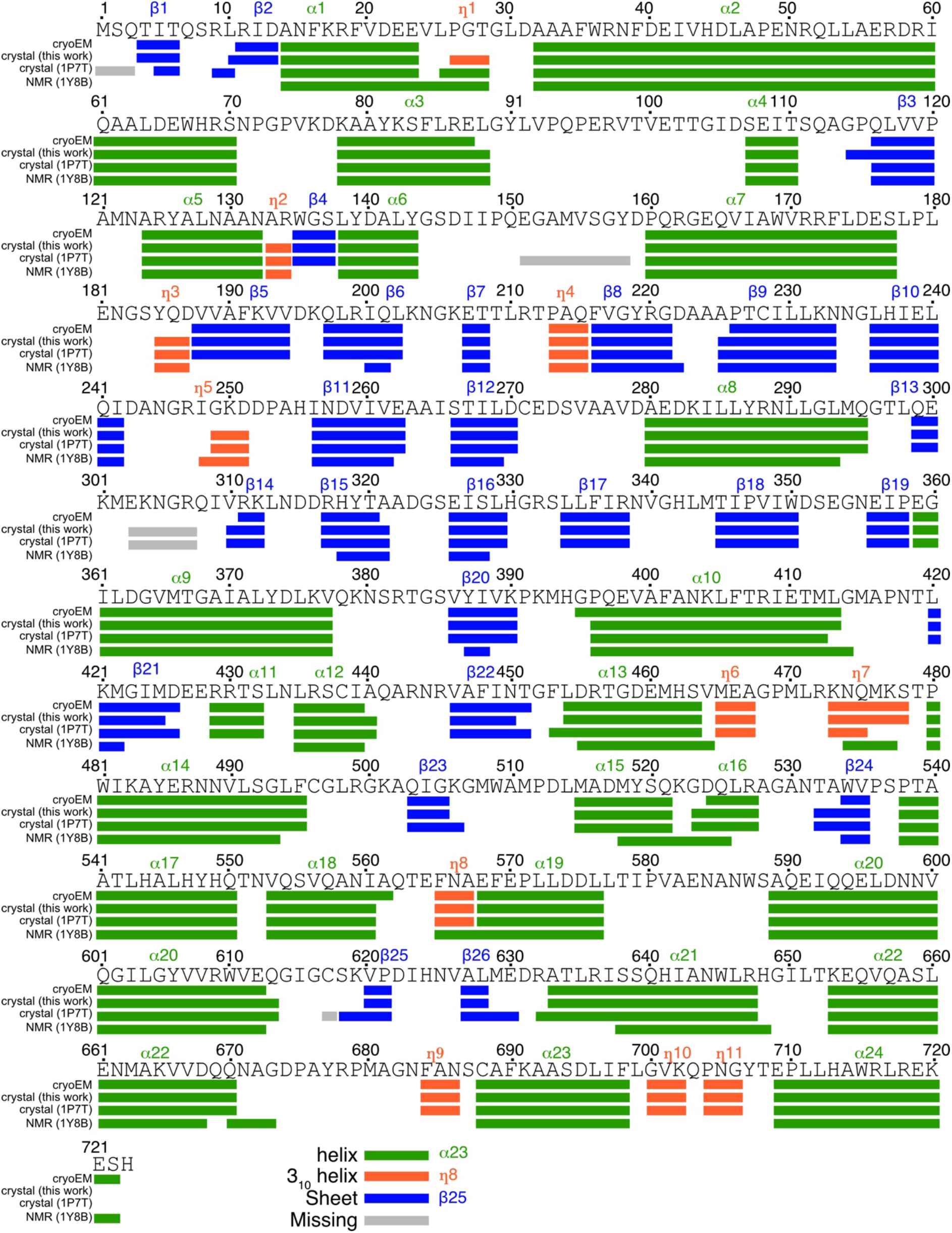
Analysis of the secondary structures of four ecMSG structures. The analysis was conducted in ENDscript ^42^. α-helices (symbol: α), 3_10_-helices (symbol: η), and β-strands (symbol: β) have been colored green, orange, and blue, respectively. Residues missing from the crystal structures are depicted by pale gray boxes.

We detected structural discrepancies between the cryo-EM and crystal structures located at loop145-161 in the α/β domain and for the 297-315 β-hairpin (Figure 2D–E). The β-hairpin 297-315 is not fully resolved in the 1.6-Å crystallographic density, as residues 303-307 are missing, an outcome consistent with previous MSG complex structures ^19,20^ and the dynamic variations revealed in the NMR ensemble. A previous NMR T_1ρ_ experiment ^24^ highlighted chemical exchanges for these two regions, implying variable conformations that inhibit structural observation. Consequently, we adopted an MSG structure predicted using Alphafold2 ^25,26^ (UniProt ID: P37330) as a reference for the β-hairpin region. Apo-form ecMSG is extensively similar to the MSG:pyruvate:acetyl-CoA complex structure (RMSD 0.74 Å). In inspecting the ecMSG catalytic pocket of all available structures, we noted that most residues involved in binding to acetyl-CoA and pyruvate are shared across methodologies, except for F453, L454, M629, and E630 (Figure 2E). Sidechains of M629 and E630 are strikingly oriented out from the catalytic core, thereby increasing the exposed surface hosting the acetyl carbon chain of acetyl-CoA. A flipped F453 aromatic ring and displacement of the L454 sidechain toward the catalytic core also tightly clamp the substrate (pyruvate, in the case of 1P7T), resulting in a stable MSG:pyruvate:acetyl-CoA complex.

Structural variations among GroEL oligomers have been presented previously based on cryo-EM ^27^, highlighting dynamic features of the protein under the snapshot frozen state. Recently developed mathematical algorithms for cryo-EM analyses— including 3D variability analysis (3DVA) ^28^, cryoDRGN ^29^ and multi-body refinement ^30^—have enabled further elucidation of the local structural dynamics of reconstructed maps. Thus, it is now feasible to compare the conformational changes of ecMSG calculated from the cryo-EM map, the atomic displacement parameters (ADPs, also known as B factors) obtained from crystal structures, and from NMR-derived S^2^ order parameters. The crystal structure of ecMSG exhibits ADPs of 20-30 across the entire protein, with high ADPs attributable to four residue clusters (132-162, 305-325, 580-590 and 670-685) notably associated with the α/β domain (Figure 4A). Application of 3DVA further revealed continuous structural changes in the ecMSG cryo-EM map. For instance, 3 of the 20 3DVA-derived frames shown in Figure 4B highlight conformational heterogeneities in ecMSG, primarily attributable to the α/β domain and β-hairpin. Thus, cryo-EM variabilities and crystallographic ADPs both consistently portray the structural dynamics of ecMSG. Moreover, NMR-measured spin relaxation and S^2^ order parameters derived from published ^13^C chemical shifts (BMRB ID: 5471) ^31^ (http://www.randomcoilindex.ca/) also describe these ecMSG dynamics. The high NMR T_1ρ_ values of apo-ecMSG measured previously ^24^ further support the potential for conformational exchanges in the four regions unveiled by cryo-EM 3DVA and crystallographic ADPs. In an overlaid plot of crystal ADPs and reverse NMR S^2^ order parameters (1-S^2^), these four regions are strongly correlated (Figure 4C), implying that ecMSG dynamics in solution represented by both NMR T_1ρ_ and S^2^ values are consistent among all three structural biology tools.

**Figure 4.**
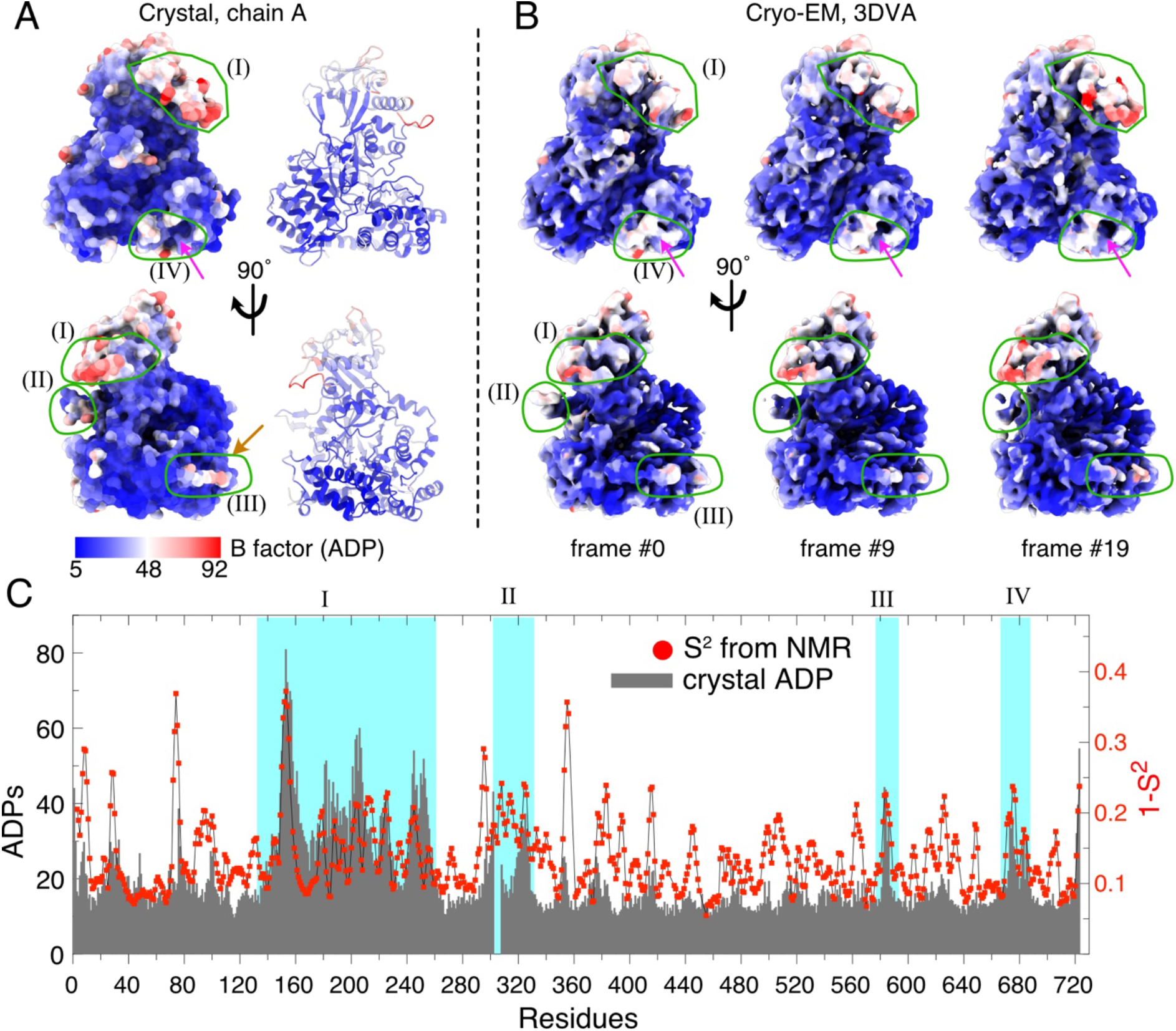
Structural dynamics of ecMSG. (A) The crystal structure of apo-form ecMSG is presented in surface or cartoon format colored according to B factors. The four circled regions(I, II, III and IV) denote local clusters with high B factor values. (B) The cryo-EM map of ecMSG was analyzed by means of 3D variability analysis (3DVA). Three frames were selected to represent the structural heterogeneity of the four regions circled in (A). The maps are colored according to the ADP values from the crystal structure of ecMSG to indicate correlation between cryo-EM map variations and ADPs in the crystal structure. The magenta arrows indicate structural changes in region IV. (C) The residue-specific ADPs and reverse NMR S^2^ (1-S^2^) values of apo-form ecMSG. The four regions circled in (A) and (B) are highlighted in cyan to reveal strong correlation between NMR dynamics and crystal stabilities.

### The cryo-EM structure of a dimeric MSG

While processing the ecMSG cryo-EM data, we noticed some ecMSG particles tended to cluster in the micrographs, resulting in a dimeric form. We deployed a large box (260 Å) for 2D classification of this dimeric ecMSG, as shown in Figure 5A. The structural features of the ecMSG dimer in the 2D classes are consistent with its monomeric form (Figure 5B), indicating that: (1) the dimeric form physically interacts in a non-random manner; and (2) there was an opportunity to obtain a 3D map of dimeric ecMSG. We targeted 15,832 particles (6% of all 250K MSG particles) of clear dimeric ecMSG to reconstruct a 4.14Å map. The infrequent dimer certainly explains its invisibility in the SEC-MALS data (Figure 1A), as the UV or particle-scattering signals are as weak as background noise. Dimeric MSG from *Mycobacterium tuberculosis* (mtMSG) has been reported previously ^32^, but its frequency is not as low as we have characterized here from cryo-EM of ecMSG. Although the map resolution was limited by the number of particles available to us and their distributed orientations, we could clearly place two MSG monomers into the map without structural rearrangements, evidencing an asymmetric and ordered assembly. The interfaces between the two ecMSG monomers differ from those currently known from ecMSG and mtMSG crystal structures, implying a novel dimeric structure under solution-like conditions. Our cryo-EM map reveals that the dimeric ecMSG is assembled via the α/β-domain of one ecMSG interacting with that of the other monomer via electrostatic interactions (Figure 5C). Moreover, the β-hairpins (residues 297-315) of both monomers are in close proximity. Due to the limited resolution and dynamic features of the β-hairpin, our cryo-EM map and docked model could not resolve additional features of this interaction. As the catalytic core pockets of the individual ecMSG monomers are not blocked in the dimeric state, we postulate that the dimer is not auto-inhibited or inactive. Accordingly, the biological role, if any, of this dimeric form warrants further analysis.

**Figure 5.**
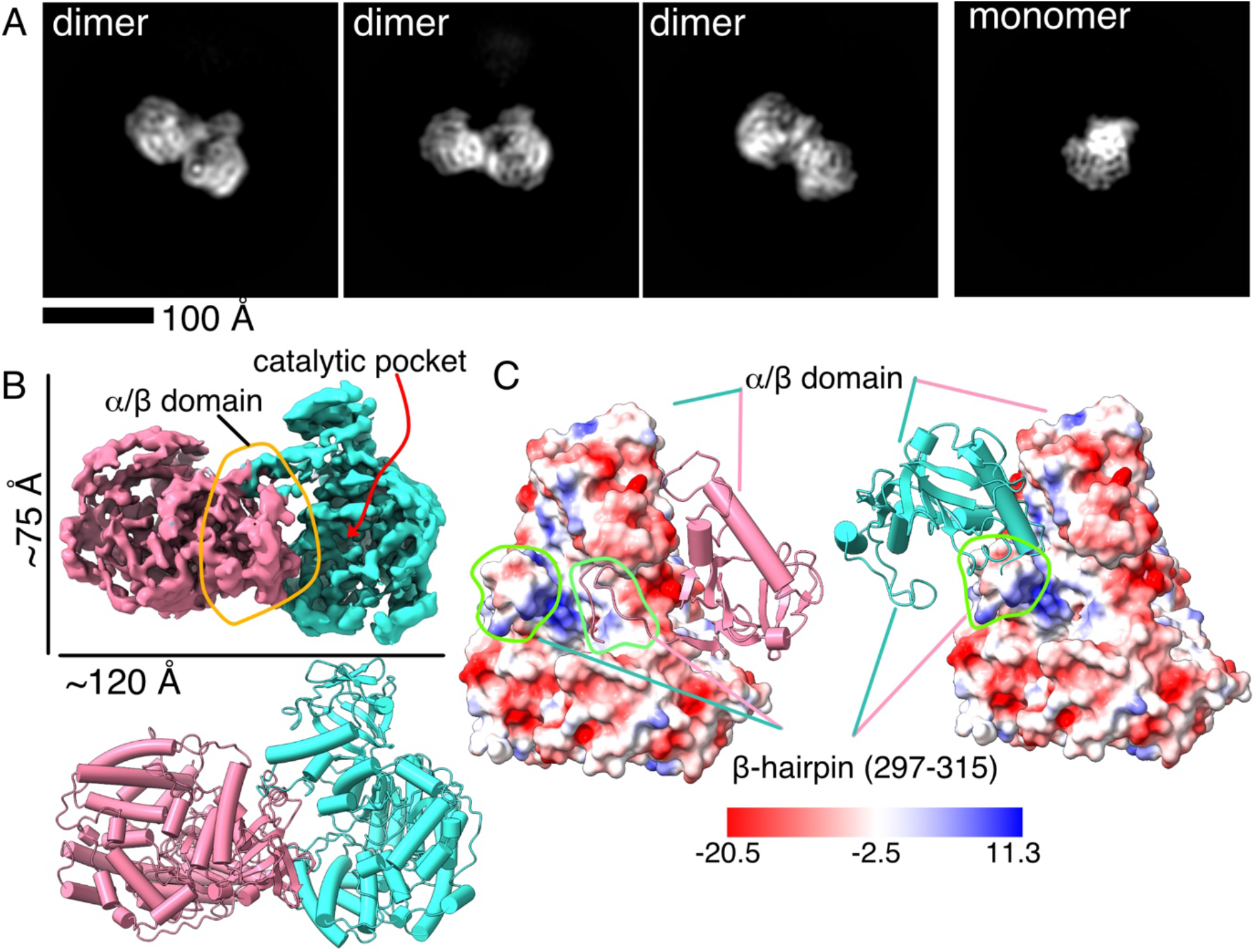
Dimeric ecMSG structure. (A) Representative 2D classification images of dimeric ecMSG using a 263Å×263Å box. For comparison, a monomeric form is also shown (rightmost panel). The interacting ecMSG monomers present identical signals and contrast. The particle size of dimeric ecMSG is ~100 Å, i.e., wider than the referenced monomeric form (~70 Å). (B) Reconstructed dimeric ecMSG map. The two ecMSG molecules, shown in salmon and cyan, are well aligned on the colored map. Notably, the catalytic pocket is not spatially hindered in this dimeric form. (C) Two views of dimeric ecMSG are shown in surface-charge and cartoon formats to highlight inter-chain interactions. The α/β domain of one ecMSG monomer interacts with the lateral side of its paired ecMSG monomer, mainly via electrostatic interactions. The two dynamic β-hairpins comprising residues 297-315 from each monomer have been labeled and circled, revealing their spatial proximity in the dimeric form.

### Conclusion

In this work, we have demonstrated that the structure of 723-residue ecMCG can be well determined by cryo-EM technology. The overall structures and details of the residue sidechains of the protein structures resolved by both cryo-EM and X-ray crystallography are significantly identical, implying that the cryo-EM structure of ecMSG we present herein is reliable. Our high-quality ecMSG cryo-EM structure validates the feasibility of routinely deploying cryo-EM for structural determinations of sub-100 kDa proteins. Dynamic ecMSG variabilities derived from cryo-EM analysis, crystallographic ADPs and NMR order parameters are highly consistent, evidencing the reliability of structural heterogeneities illustrated by cryo-EM. Significantly, investigations of mtMSG using crystallography to identify inhibitors necessitated co-crystallization of the protein with its ligand ^16,17^, thus impeding the efficiency of drug discovery. Our demonstration here of the feasibility of cryo-EM-based structural studies of ligand-bound MSG may resolve this impediment.

It is worth noting our ecMSG cryo-EM dataset is relatively small (700 micrographs, ~211,000 particles) for an asymmetric protein. Recently developed direct detectors such as K3 (Gatan), Apollo (Direct Electron), and Falcon IV (Thermo), accompanied with super-resolution ^33^ and electron-event representation (EER) ^34^ data collection modes, could further enhance the quality of micrographs and particle collection. Finally, a high-quality specimen using an Au-specific hexagonal grid ^35^ and a large dataset may also improve map resolution and reveal details of protein sidechains. Thus, we have demonstrated that cryo-EM can be deployed to characterize ligand- or drug-bound structures of sub-100 kDa protein-ligand/drug structures.

## Methods

### Protein production

The *MSG* gene was amplified directly from the *E. coli* DH5**α** genome and sub-cloned into the pRSFDuet-1 (Millipore) expression vector, followed by an N-terminal MGSSHHHHHHGSSGENLYFQGS (His-tev) sequence that encodes a His-tag and a sequence for tobacco etch virus (TEV) protease. Plasmid hosting the His-tev-ecMSG construct was transformed into *E. coli* BL21 DE3 strain. Two liters of cultured cells was induced by 0.6 mM IPTG at 20 °C for 16 h, until the optical density (OD_600_) reached 0.8. After sonication and centrifugation, His-tagged ecMSG was loaded onto a gravity nickel affinity column (Roche) and eluted using 300 mM imidazole. The His-tev tag was retained for samples subjected to cryo-EM and SEC-MALS, but it was cleaved by means of TEV protease treatment at 4 °C overnight for crystallization experiments. The resulting ecMSG proteins were then concentrated and subjected to size exclusion chromatography for further purification. The proteins were switched to buffer comprising 25 mM Tris pH 7.6, 200 mM NaCl and 4 mM β-mercaptoethanol using an Akta FPLC system (GE Healthcare). Pure His-tev-ecMSG or tag-cleaved ecMSG was concentrated to 40-80 mg/ml. Aliquoted ecMSG proteins were frozen in liquid nitrogen and stored at −80 °C.

### Size exclusion chromatography combined with multi-angle light scattering (SEC-MLAS)

SEC-MALS measurements were conducted using an Agilent 1260 Infinity HPLC system equipped with a Wyatt miniDAWN TREOS detector and an Optilab T-rEX differential refractive index detector (Wyatt). Bovine serum albumin (A1900, Sigma) was used for system calibration. The SEC experiments were performed on an Agilent Bio SEC-3 300Å 4.6×300 mm system equilibrated in a buffer containing 25 mM Tris-HCl pH 7.6, 200 mM NaCl and 0.02% NaN_3_. His-tev-ecMSG (100 μg) was injected into the column at a flow rate of 0.35 ml/min. The theoretical molecular weight of His-tev-MSG is 82.9 kDa. The experimental molecular weight of His-tev-ecMSG was calculated using ASTRA6 software (Wyatt), with a d*n*/d*c* value of 0.185 ml/g.

### Cryo-EM data collection, processing, and model building

Quantifoil grids (200 mesh Cu, R2.0/1.0) were glow-discharged for 15 sec before preparing ecMSG samples for cryo-EM. Sample specimens were generated using Vitrobot Mark IV (Thermo Fischer Scientific) with 4 μl of 0.3-1.0 mg/ml ecMSG proteins and a blotting time of 4 sec at 100% humidity and 4 °C. The blotted grids were frozen in nitrogen-cooled liquid ethane. The cryo-EM data for ecMSG was collected on a Titan Krios electron microscope, operated at 300 keV, and equipped with a K2 GIF-Quantum direct electron detector (Gatan). The slit width was 20 eV and the microscope nominal magnification of 165,000 resulted in a calibrated physical pixel size of 0.822 Å/pixel. The gain reference was collected before data collection. Data was acquired using Thermo EPU 2.1. We used 80 frames of a total electron dose of 51.2 (e-/Å^2^) to acquire 700 micrographs of ecMSG, with a defocus range of −1.0 to −1.8 μm. The statistics for cryo-EM-reconstructed maps, as well as a built structure, are listed in Table S1.

The cryo-EM dataset was processed in cryoSPARC (3.2) ^23^, with an illustrated workflow shown in Figure S1. All movie frames were used for alignment using the Patch Motion tool in cryoSPARC. Contrast transfer function parameters were estimated using patchCTF. We used 50 aligned micrographs to select ecMSG particles based on a blob diameter of 65-90 Å. Selected particles were extracted and grouped according to 2D classification, and then three ecMSG-containing 2D classes were selected for template-based particle picking from all ecMSG micrographs. We extracted a total of 584,461 particles from 700 dose-weighted micrographs. Further polishing by 2D classification resulted in 254,442 particles. We generated three *ab initio* models of ecMSG in cryoSPARC for heterogeneous 3D classification. We selected 211,582 particles for further homogeneous refinement, followed by CTF refinement and non-uniform sampling refinement. The gold standard resolution (FSC_0.143_) of the final ecMSG map using the combined datasets is 2.89 Å. An analysis of local resolution estimation of this 2.89-Å ecMSG map was conducted in cryoSPARC (Figure S1). Additionally, a 3D variability analysis (3DVA) ^28^ of the ecMSG particles was performed with 3 modes and filtered at 6 Å, with the display in linear mode revealing the structural heterogeneity of ecMSG.

Model building of ecMSG using the 2.89-Å map was performed in Phenix (1.19) ^36^ using previous crystal MSG structures without the ligands pyruvate and acetyl-CoA. The model was first refined using real-space refinement with non-crystallographic symmetry (NCS) and secondary structure restraints. Coot 0.9.1 ^37^ was used to inspect, build and modify ecMSG residues manually. ISOLDE 1.3 ^38^ was deployed to fix rotameric outliers and steric clashes, resulting in an ecMSG structure of satisfactory quality. The ecMSG structure was validated using Molprobity ^39^, giving rise to a clash score of 1.69. Model quality is summarized in Table S1.

## Crystallography

Sitting drop crystal screenings of 38 mg/ml apo-form ecMSG were performed using Phoenix (Rigaku), and the initial hit was found after 7 days in buffer containing 0.1 M Tris pH 7.5, 0.2 M NaCl, and 20% PEG3350 at 22 °C. Crystal growth was further optimized using a wide range of protein concentrations, pH, and percentages of PEG 4,000 or PEG 8,000. The optimized sitting-dropped ecMSG crystals grew within 2 weeks in a buffer of 0.1 M Tris pH 8.5, 0.2 M NaCl, and 19-22% PEG4000 at 23 °C. The ecMSG crystals were harvested upon addition of 10-15% glycerol or ethylene glycol as a cryogenic protectant for X-ray diffraction data collection at the TPS-05A beamline station of the National Synchrotron Radiation Research Center, Hsinchu, Taiwan. In total, we collected 180 diffraction images, with an oscillation frame rate of 1 degree. Diffraction data were processed and cut at 1.6 Å using HKL2000 software ^40^, resulting in space group P212121. Molecular replacement using a published MSG:pyruvate:acetyl-CoA structure (PDB ID:1P7T) ^20^ was introduced to determine the phase and as an initial building model. The ecMSG structure was iteratively built, refined, manually inspected, and polished using Phenix 1.19 and Coot 0.9.1. The final ecMSG structure was validated in Molprobity. Statistics for crystallographic data are listed in Table S2.

## Supporting information

Movie representing structural dynamics of MSG

## Data availability

The atomic coordinates of ecMSG determined by cryo-EM and crystallography have been deposited in PDB with the accession codes 7YQM and 7YQN, respectively. The cryo-EM maps of ecMSG monomer and dimer have been submitted to EMDB with accession codes 34029 and 34030, respectively.

## Acknowledgment

K.-P. W. was supported by the Ministry of Science and Technology (MOST) MOST 108-2311-B-001-016-MY3 and a Career Development Award AS-CDA-110-L03 from Academia Sinica, Taipei, Taiwan. We thank the Academia Sinica Cryo-EM center (ASCEM) for assistance with data collection and appreciate the technical services provided by the protein crystallography facility of the National Synchrotron Radiation Research Center supported by MOST, Taiwan. ASCEM is jointly supported by the Academia Core Facility and Innovative Instrument Project (AS-CFII-108-110) and Taiwan Protein Project (AS-KPQ-109-TPP2).

## Notes

### Competing Interest Statement

The authors have declared no competing interest.

## References

1 Merk, A. et al. Breaking Cryo-EM Resolution Barriers to Facilitate Drug Discovery. Cell 165, 1698–1707, doi:10.1016/j.cell.2016.05.040 (2016).

2 Kuhlbrandt, W. Biochemistry. The resolution revolution. Science 343, 1443–1444, doi:10.1126/science.1251652 (2014).

3 Cheng, Y. Single-Particle Cryo-EM at Crystallographic Resolution. Cell 161, 450–457, doi:10.1016/j.cell.2015.03.049 (2015).

4 Chiu, W. & Downing, K. H. Editorial overview: Cryo Electron Microscopy: Exciting advances in CryoEM Herald a new era in structural biology. Curr Opin Struct Biol 46, iv–viii, doi:10.1016/j.sbi.2017.07.006 (2017).

5 Sprangers, R. & Kay, L. E. Quantitative dynamics and binding studies of the 20S proteasome by NMR. Nature 445, 618–622, doi:10.1038/nature05512 (2007).

6 Tugarinov, V., Hwang, P. M. & Kay, L. E. Nuclear magnetic resonance spectroscopy of high-molecular-weight proteins. Annu Rev Biochem 73,107–146, doi:10.1146/annurev.biochem.73.011303.074004 (2004).

7 Tugarinov, V. & Kay, L. E. Ile, Leu, and Val methyl assignments of the 723-residue malate synthase G using a new labeling strategy and novel NMR methods. J Am Chem Soc 125, 13868–13878, doi:10.1021/ja030345s (2003).

8 Khoshouei, M., Radjainia, M., Baumeister, W. & Danev, R. Cryo-EM structure of haemoglobin at 3.2 A determined with the Volta phase plate. Nat Commun 8, 16099, doi:10.1038/ncomms16099 (2017).

9 Herzik, M. A., Jr., Wu, M. & Lander, G. C. High-resolution structure determination of sub-100 kDa complexes using conventional cryo-EM. Nat Commun 10, 1032, doi:10.1038/s41467-019-08991-8 (2019).

10 Fan, X. et al. Single particle cryo-EM reconstruction of 52 kDa streptavidin at 3.2 Angstrom resolution. Nat Commun 10, 2386, doi:10.1038/s41467-019-10368-w (2019).

11 Greber, B. J. et al. The cryoelectron microscopy structure of the human CDK-activating kinase. Proc Natl Acad Sci U S A 117, 22849–22857, doi:10.1073/pnas.2009627117 (2020).

12 Watson, Z. L. et al. Structure of the bacterial ribosome at 2 A resolution. Elife 9, doi:10.7554/eLife.60482 (2020).

13 Dong, Y. et al. Cryo-EM structures and dynamics of substrate-engaged human 26S proteasome. Nature 565, 49–55, doi:10.1038/s41586-018-0736-4 (2019).

14 Puckett, S. et al. Glyoxylate detoxification is an essential function of malate synthase required for carbon assimilation in Mycobacterium tuberculosis. Proc Natl Acad Sci U S A 114, E2225–E2232, doi:10.1073/pnas.1617655114 (2017).

15 Anstrom, D. M. & Remington, S. J. The product complex of M. tuberculosis malate synthase revisited. Protein Sci 15, 2002–2007, doi:10.1110/ps.062300206 (2006).

16 Smith, C. V. et al. Biochemical and structural studies of malate synthase from Mycobacterium tuberculosis. J Biol Chem 278, 1735–1743, doi:10.1074/jbc.M209248200 (2003).

17 Huang, H. L., Krieger, I. V., Parai, M. K., Gawandi, V. B. & Sacchettini, J. C. Mycobacterium tuberculosis Malate Synthase Structures with Fragments Reveal a Portal for Substrate/Product Exchange. J Biol Chem 291, 27421–27432, doi:10.1074/jbc.M116.750877 (2016).

18 McVey, A. C. et al. Structural and Functional Characterization of Malate Synthase G from Opportunistic Pathogen Pseudomonas aeruginosa. Biochemistry 56, 5539–5549, doi:10.1021/acs.biochem.7b00852 (2017).

19 Howard, B. R., Endrizzi, J. A. & Remington, S. J. Crystal structure of Escherichia coli malate synthase G complexed with magnesium and glyoxylate at 2.0 A resolution: mechanistic implications. Biochemistry 39, 3156–3168, doi:10.1021/bi992519h (2000).

20 Anstrom, D. M., Kallio, K. & Remington, S. J. Structure of the Escherichia coli malate synthase G:pyruvate:acetyl-coenzyme A abortive ternary complex at 1.95 A resolution. Protein Sci 12, 1822–1832, doi:10.1110/ps.03174303 (2003).

21 Grishaev, A., Tugarinov, V., Kay, L. E., Trewhella, J. & Bax, A. Refined solution structure of the 82-kDa enzyme malate synthase G from joint NMR and synchrotron SAXS restraints. J Biomol NMR 40, 95–106, doi:10.1007/s10858-007-9211-5 (2008).

22 Tugarinov, V., Choy, W. Y., Orekhov, V. Y. & Kay, L. E. Solution NMR-derived global fold of a monomeric 82-kDa enzyme. Proc Natl Acad Sci U S A 102, 622–627, doi:10.1073/pnas.0407792102 (2005).

23 Punjani, A., Rubinstein, J. L., Fleet, D. J. & Brubaker, M. A. cryoSPARC: algorithms for rapid unsupervised cryo-EM structure determination. Nat Methods 14, 290–296, doi:10.1038/nmeth.4169 (2017).

24 Tugarinov, V. & Kay, L. E. Quantitative NMR studies of high molecular weight proteins: application to domain orientation and ligand binding in the 723 residue enzyme malate synthase G. J Mol Biol 327, 1121–1133, doi:10.1016/s0022-2836(03)00238-9 (2003).

25 Jumper, J. et al. Highly accurate protein structure prediction with AlphaFold. Nature 596, 583–589, doi:10.1038/s41586-021-03819-2 (2021).

26 Jumper, J. & Hassabis, D. Protein structure predictions to atomic accuracy with AlphaFold. Nat Methods 19, 11–12, doi:10.1038/s41592-021-01362-6 (2022).

27 Roh, S. H. et al. Subunit conformational variation within individual GroEL oligomers resolved by Cryo-EM. Proc Natl Acad Sci U S A 114, 8259–8264, doi:10.1073/pnas.1704725114 (2017).

28 Punjani, A. & Fleet, D. J. 3D variability analysis: Resolving continuous flexibility and discrete heterogeneity from single particle cryo-EM. J Struct Biol 213, 107702, doi:10.1016/j.jsb.2021.107702 (2021).

29 Zhong, E. D., Bepler, T., Berger, B. & Davis, J. H. CryoDRGN: reconstruction of heterogeneous cryo-EM structures using neural networks. Nat Methods 18, 176–185, doi:10.1038/s41592-020-01049-4 (2021).

30 Nakane, T., Kimanius, D., Lindahl, E. & Scheres, S. H. Characterisation of molecular motions in cryo-EM single-particle data by multi-body refinement in RELION. Elife 7, doi:10.7554/eLife.36861 (2018).

31 Berjanskii, M. V. & Wishart, D. S. A simple method to predict protein flexibility using secondary chemical shifts. J Am Chem Soc 127, 14970–14971, doi:10.1021/ja054842f (2005).

32 Kumar, R. & Bhakuni, V. A functionally active dimer of mycobacterium tuberculosis malate synthase G. Eur Biophys J 39, 1557–1562, doi:10.1007/s00249-010-0598-7 (2010).

33 Feathers, J. R., Spoth, K. A. & Fromme, J. C. Experimental evaluation of super-resolution imaging and magnification choice in single-particle cryo-EM. J Struct Biol X 5, 100047, doi:10.1016/j.yjsbx.2021.100047 (2021).

34 Guo, H. et al. Electron-event representation data enable efficient cryoEM file storage with full preservation of spatial and temporal resolution. IUCrJ 7, 860–869, doi:10.1107/S205225252000929X (2020).

35 Naydenova, K., Jia, P. & Russo, C. J. Cryo-EM with sub-1 A specimen movement. Science 370, 223–226, doi:10.1126/science.abb7927 (2020).

36 Liebschner, D. et al. Macromolecular structure determination using X-rays, neutrons and electrons: recent developments in Phenix. Acta Crystallogr D Struct Biol 75, 861–877, doi:10.1107/S2059798319011471 (2019).

37 Casanal, A., Lohkamp, B. & Emsley, P. Current developments in Coot for macromolecular model building of Electron Cryo-microscopy and Crystallographic Data. Protein Sci 29, 1069–1078, doi:10.1002/pro.3791 (2020).

38 Croll, T. I. ISOLDE: a physically realistic environment for model building into low-resolution electron-density maps. Acta Crystallogr D Struct Biol 74, 519–530, doi:10.1107/S2059798318002425 (2018).

39 Prisant, M. G., Williams, C. J., Chen, V. B., Richardson, J. S. & Richardson, D. C. New tools in MolProbity validation: CaBLAM for CryoEM backbone, UnDowser to rethink “waters,” and NGL Viewer to recapture online 3D graphics. Protein Sci 29, 315–329, doi:10.1002/pro.3786 (2020).

40 Otwinowski, Z. & Minor, W. Processing of X-ray diffraction data collected in oscillation mode. Methods Enzymol 276, 307–326 (1997).

41 Pettersen, E. F. et al. UCSF ChimeraX: Structure visualization for researchers, educators, and developers. Protein Sci 30, 70–82, doi:10.1002/pro.3943 (2021).

42 Robert, X. & Gouet, P. Deciphering key features in protein structures with the new ENDscript server. Nucleic Acids Res 42, W320–324, doi:10.1093/nar/gku316 (2014).

