## Supporting information for "Cryo-EM reveals the structure and dynamics of a 723-residue malate synthase G"

Figure S1 Workflow of cryo-EM data processing.

Table S1 Cryo-EM data collection, refinement, and validation statistics for ecMSG.

Table S2 Crystal data collection, refinement and validation statistics for ecMSG.

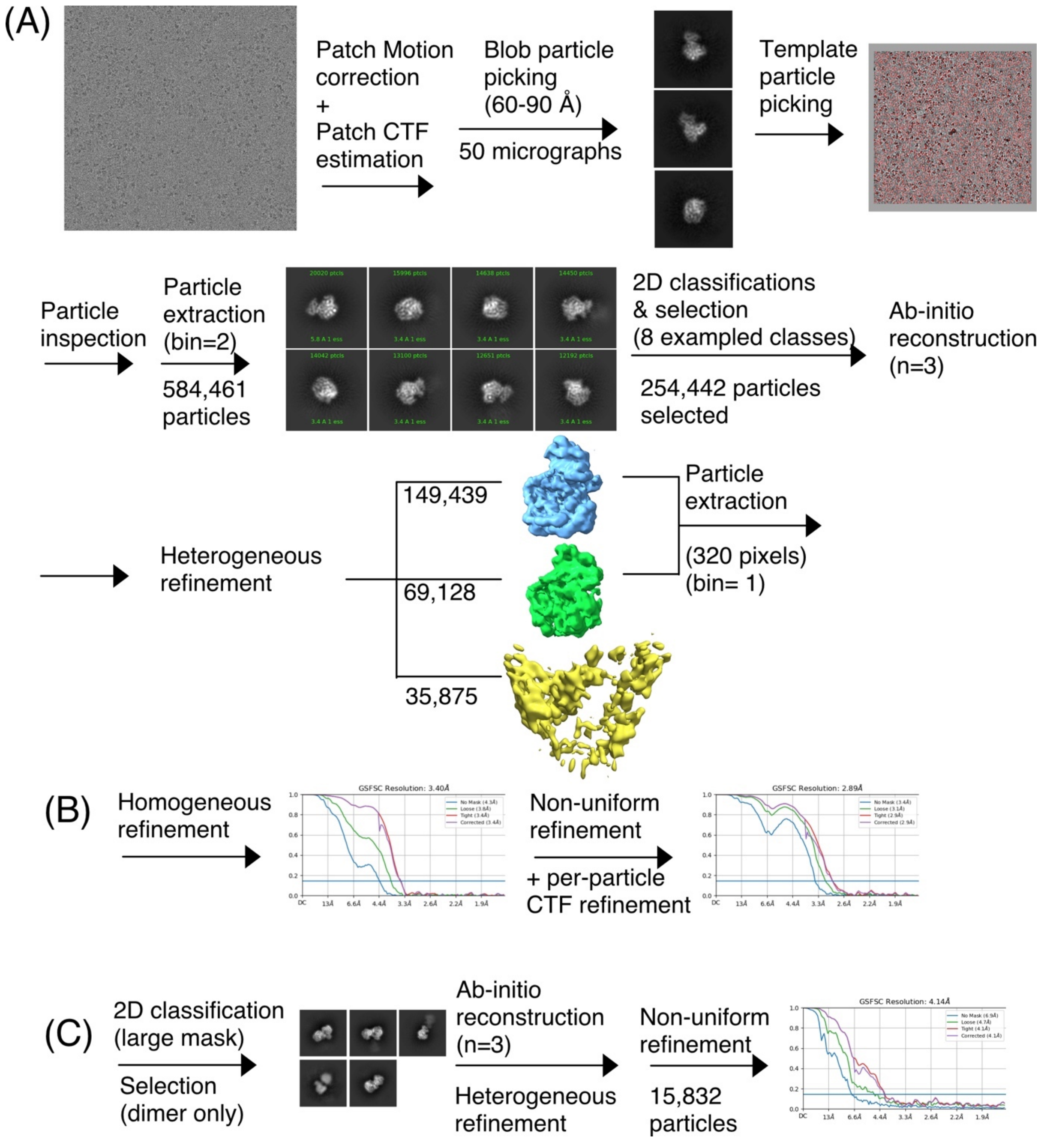

Figure S1. Workflow for cryo-EM data processing. All processes were conducted using cryoSPARC 3.2. (A) Steps from motion correction, CTF estimation, particle picking, 2D classifications, selection of 2D classes, ab-initio model reconstruction, to 3D classifications are presented. (B) Appropriate 3D classes were selected to perform 3D refinements, including homogeneous and non-uniform refinements. The map resolution is 2.89 Å. (C) The same appropriate 3D classes used in (B) were further subjected to 2D classification to select clear dimeric groups for 3D map reconstruction. The final resolution is 4.14 Å.

**Table 1. Cryo-EM data collection, refinement, and validation statistics for ecMSG.**

| | Monomer | Dimer |
|---|---|---|
| **Data collection and processing** | | |
| Microscope/detector | Krios/K2 | |
| Magnification | 165,000 | |
| Voltage | 300 kV | |
| Electron exposure (e-/Å$^2$) | 51.2 | |
| Defocus range (μm) | -1.0 to -1.8 | |
| Pixel size (Å) | 0.822 | |
| Symmetry imposed | C1 | |
| Initial particle images (no.) | 584,461 | |
| Final particle images (no.) | 211,582 | 15,832 |
| Reconstruction method | Single particle reconstruction | |
| Map resolution (Å) ($FSC_{0.143}$) | 2.89 | 4.14 |
| Map sharpening B factor (Å$^2$) | 94.4 | 77.2 |
| **Refinement** | | |
| Initial model | 1P7T | EM monomer |
| **Model composition** | | |
| Non-hydrogen atoms | 5656 | |
| Protein residues | 723 | |
| Ligand / Water | 0 / 0 | |
| **B factor (Å$^2$) (min / max / mean)** | | |
| Protein | 25.80 / 61.84 / 29.09 | |
| **R.m.s. deviations** | | |
| Bond lengths (Å) | 0.004 | |
| Bond angles (˚) | 0.581 | |
| **Validation** | | |
| MolProbity score | 1.25 | |
| Clash score | 1.69 | |
| Poor rotamers (%) | 0.17 | |
| **Ramachandran plot** | | |
| Favored (%) | 95.28 | |
| Allowed (%) | 4.72 | |
| Outliers (%) | 0 | |
| **Model vs Data** | | |
| CC (mask) | 0.78 | |
| dFSC model (0 / 0.143 / 0.5) (Å) | 2.8 / 2.8 / 3.1 | |
| Accession code (EMDB) | 34029 | 34030 |
| Accession code (PDB) | 7YQM | |

**Table 2. Crystal data collection, refinement and validation statistics for ecMSG.**

| **Data collection** | |
|---|---|
| Source | NSRRC TPS05A |
| Wavelength (Å) | 1.0000 |
| Space group | $P2_12_12_1$ |
| Cell dimensions | |
| a, b, c (Å) | 98.94, 101.25, 170.24 |
| α, β, γ (°) | 90.0, 90.0, 90.0 |
| Resolution (Å) | 50-1.60 (1.66-1.60) |
| Completeness (%) | 98.48 (96.78) |
| Total reflections | 221574 (21629) |
| Unique reflections | 221324 (21558) |
| Wilson B-factor | 17.75 |
| Redundancy | 4.9 (3.9) |
| $R_{merge}$ (%) | 6.7 (70.0) |
| $< I >/\sigma(I)$ | 23.30 (2.00) |
| **Refinement** | |
| Resolution (Å) | 27.45-1.60 (1.657-1.60) |
| Reflections (work/free) | 221309 (21554) |
| $R_{work}$ (%) | 15.63 (23.59) |
| $R_{free}$ (%) | 17.25 (23.84) |
| Number of atoms | 13174 |
| Protein | 11255 |
| Ligand | 38 |
| Water | 1881 |
| Average *B*-factors (Å$^2$) | 24.82 |
| Protein | 22.69 |
| Ligand | 43.12 |
| Water | 37.21 |
| RMSD | |
| Bond lengths (Å) | 0.006 |
| Bond angles (°) | 0.835 |
| MolProbity | |
| Favored (%) | 98.80 |
| Allowed (%) | 1.20 |
| Outliers (%) | 0.00 |
| Clash score | 1.33 |
| MolProbity score | 0.86 |
| PDB code | 7YQN |

*Values in parentheses are for the highest resolution shell.
